# DiffTopo: Fold exploration using coarse grained protein topology representations

**DOI:** 10.1101/2024.02.01.578456

**Authors:** Yangyang Miao, Bruno Correia

**Affiliations:** STI, EPFL, Lausanne, 1015, Swizerland

**Author notes:** {,}.

## Abstract

A major challenge in the field of computational de novo protein design is the exploration of uncharted areas within protein structural space, i.e., generating “designable” protein structures that nature has not explored. However, the large degrees of freedom of protein structural backbones complicate the sampling process during protein design. In this work, we propose a new coarse grained protein structure representation method DiffTopo - an E(3) Equivariant 3D conditional diffusion model, which greatly increases the sampling efficiency. Combined with the RFdiffusion framework, novel protein folds can be generated rapidly, allowing for efficient exploration of the designable topology space. This opens up possibilities to solve the problem of generating new folds as well to functionalize de novo proteins through motif scaffolding, where functional or enzymatic sites can be introduced into novel protein frameworks.

## 1 Introduction

Proteins govern vital biological functions, including enzyme catalysis, molecular transport, and cellular activity modulation. The intricate relationship between protein function and three-dimensional architecture is essential. One of the most important goals of the field of de novo protein design involves the computational generation of novel and realistic protein structures that fulfill specific structural and/or functional requirements (Pan & Kortemme, 2021; Kuhlman & Bradley, 2019).

Much work has been done to address the problem of computationally generating new protein structures, but has often encountered challenges in creating diverse and realistic folds. Traditional methods typically apply heuristics to assemble experimentally analyzed protein fragments into structures, which are limited by expert knowledge and available data (Simons et al., 1997; Minami et al., 2023; Mackenzie et al., 2016; Jacobs et al., 2016). Recently, deep generative models have been proposed to address these issues. Generative models rely on complex equivariant network architectures or loss functions to learn to generate 3D coordinates or internal torsion angles that describe protein structures (Anand et al., 2019; Anand & Achim, 2022; Lin & AlQuraishi, 2023; Luo et al., 2022; Trippe et al., 2023; Yim et al., 2023b;a; Eguchi et al., 2022; Watson et al., 2023). Equivariant architecture (Satorras et al., 2022; Jing et al., 2021; Fuchs et al., 2020) ensures that the probability density of protein structure sampling remains constant under translation and rotation. RFdiffusion (Watson et al., 2023) and Framediff (Yim et al., 2023b) have successfully learned the complex distribution of protein backbones and can generate designable protein backbones. However, these methods still encounter some obstacles in the goal of generating novel, previously unseen structures. For example, when generating backbones with a length less than 200 amino acids (AA), the resulting backbones are very similar to natural structures. The reason is when the length of the amino acid increases, the degree of freedom of the backbone increases exponentially. This is addressed in the work of Taylor et al. (2008) and Harteveld et al. (2022); Yang et al. (2020) where protein structure is conceptualized as a spatial stacking of standard secondary structures, significantly reducing the degree of freedom in the protein fold space.

Motivated by recent advances, our motivation was to investigate the feasibility of utilizing deep learning models to explore the protein fold space condensed protein rerpesentations. In this work, we introduce a protein representation method termed ”Coarse-grained topology(CG topology)”(depicted in Figure 1 top row). We have established a pipeline (Figure 1 bottom row) that utilizes a simple secondary structure string description as conditional input to autonomously generate a plausible CG topology. Our approach involves training a diffusion model named DiffTopo, which learns the relative spatial position distribution of plausible secondary structures. From this model, we sample and generate CG topology representations with reasonable secondary structures. We then construct the standard secondary structure at the positions specified by CG topology, resulting in what we term a ”protein sketch.” This Protein Sketch serves as direct input to RFdiffusion, to generate a designable backbone consistent with the input. Notably, our method avoids direct sampling using explicit atomic representations, ensuring a fast and efficient sampling process. Difftopo’s sampling is oriented towards exploring diverse protein folds, while RFdiffusion is subsequently employed to identify the corresponding designable backbone. This dual-stage approach contributes to the effectiveness and efficiency of our sampling methodology. The detailed description of the methodology is presented in the appendix A.1.

**Figure 1.**
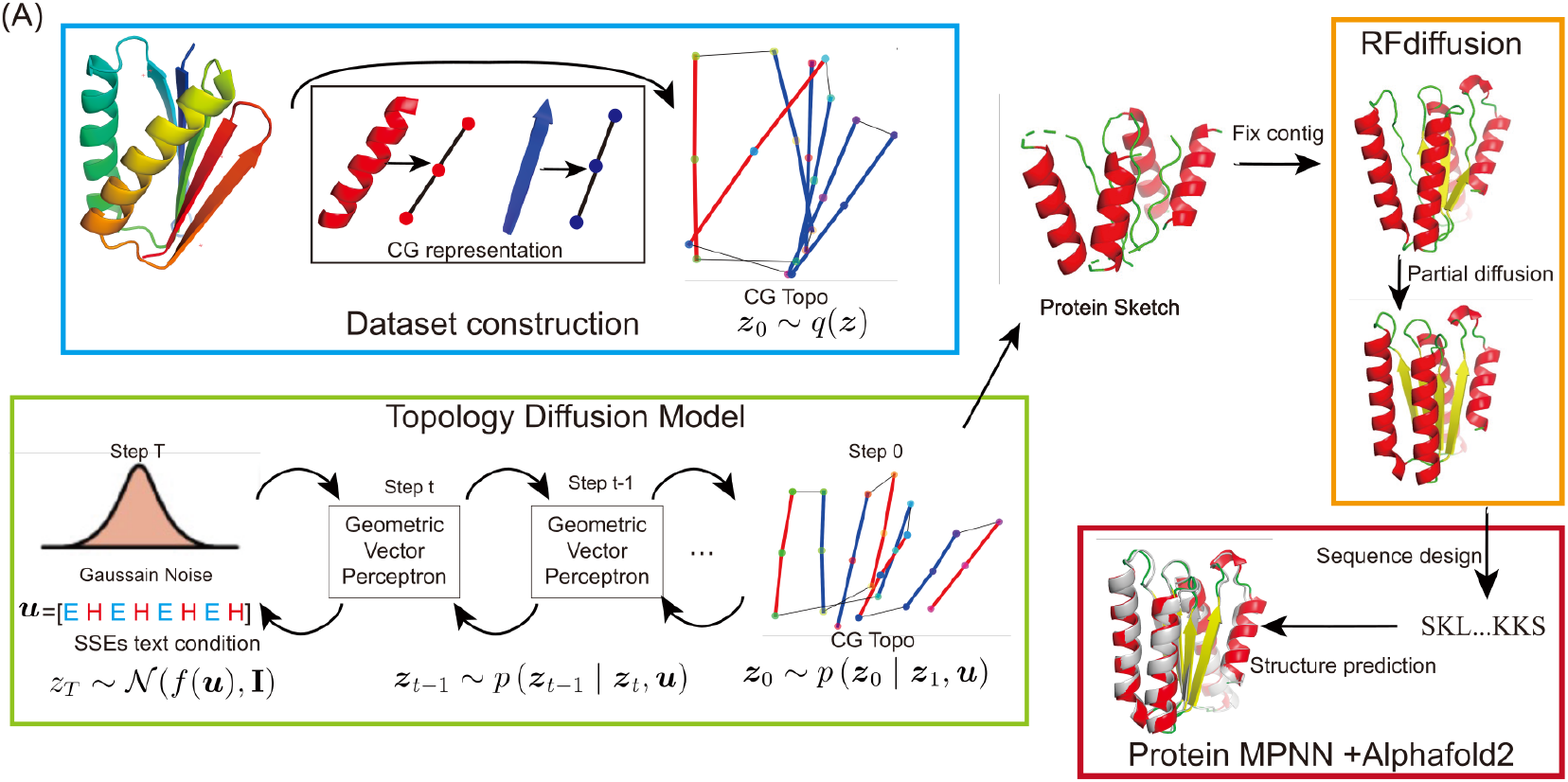
Overview of the DiffTopo de novo protein design pipeline. The first step is to convert the secondary structure into CG topology representation, followed by transforming the non-redundant backbones in the entire CATH database into CG topology to create a dataset. Then we develop the DiffTopo-RFdiffusion framework for de novo protein design. DiffTopo is the diffusion model to generate CG topology, which can be converted to protein sketches guiding RFdiffusion in the generation of protein backbones.

## 2 Results

### 2.1 DiffTopo captures the distributions of native protein structures

To assess sketch quality, standard metrics are lacking. In backbone generation tasks, evaluation often relies on geometric features like Ramachandran distribution and bond length, angle. Based on the sketch level representation, our evaluation focuses on the diffusion model’s ability to estimate data probability density, checking alignment with authentic data distribution. Figure 2A shows the selected local and pairwise geometric features used to describe CG topology data distribution. Figure 2B demonstrates the local structures of CG topology generated by diffusion model are similar with real data constructed from native backbones. Figure 2C illustrates that the relative spatial positions between CG topology also exhibit a similar distribution to that observed in real data. The results indicate that the diffusion model adequately approximates the distribution of real data, despite with a tendency to smoothen discrete high-density peaks.

**Figure 2.**
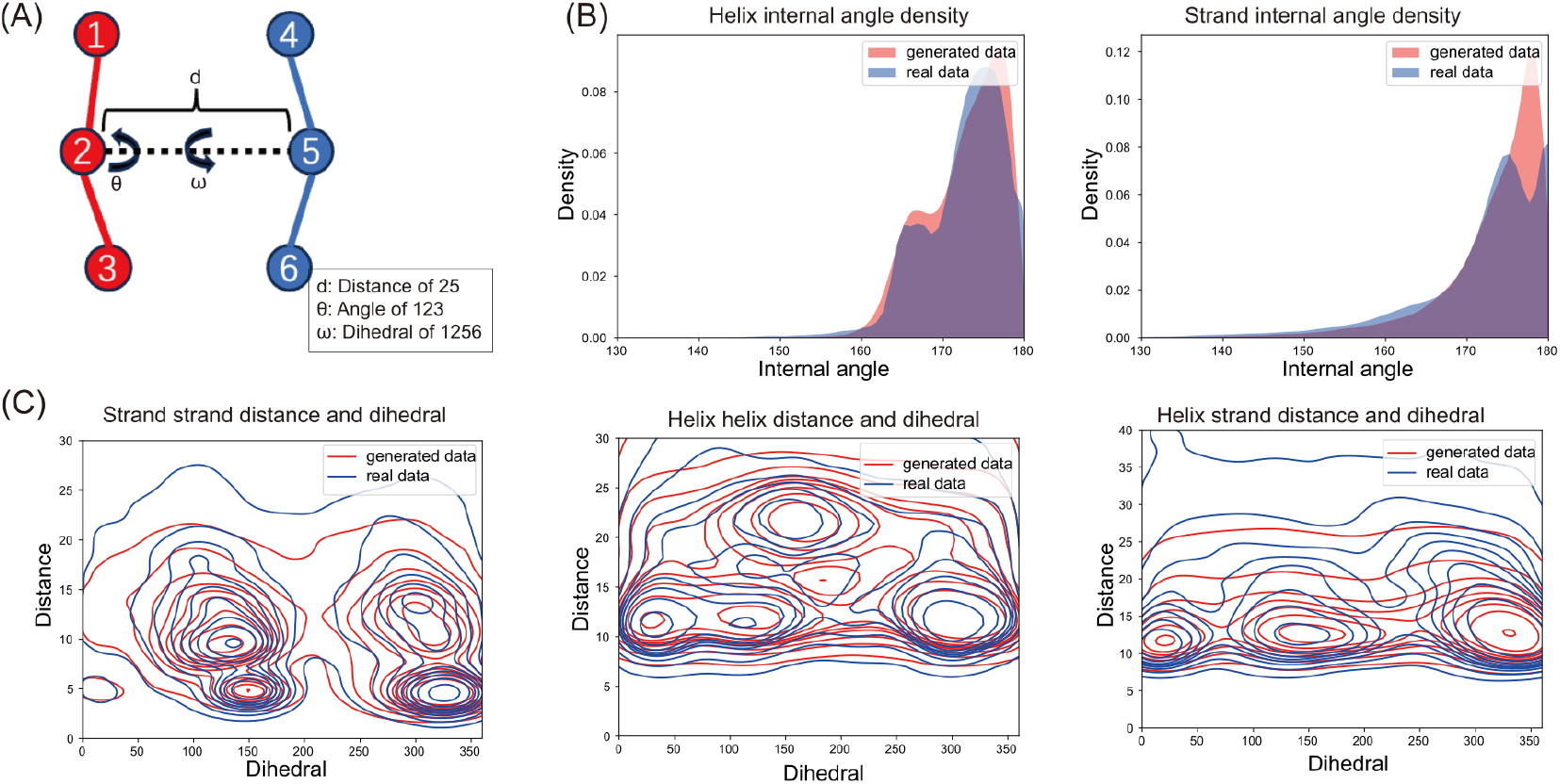
Data distribution of generated data and real data: (A) Local and pairwise geometric features to describe spatial relative positions of SSEs, including internal angle inside one SSE, distance between centroids of two SSEs and dihedral between two SSEs. (B) Local geometric feature distributions of generated (red) and real data (blue). (C) Pairwise geometric feature distributions of generated (red) and real data (blue)

### 2.2 Similarity of sketch and backbones

To confirm that the CG topology accurately represents the fold of the real protein, and to verify that the backbone generated by RFdiffusion has the same protein fold as the CG topology, we assess the similarity between protein sketches constructed from CG topology and backbones — both real and randomly generated by RFdiffusion. Here, we calculate TMscore (Zhang & Skolnick, 2004; Xu & Zhang, 2010) and RMSD metrics to compare native backbones with protein sketches derived from their CG topology. The TMscore and RMSD between sketches generated from CG topology and the corresponding backbones are also calculated. As depicted in Figure 3A and 3B, the arrangement of most Secondary Structure Elements (SSEs) in protein sketches is similar to their corresponding backbones, with TMscores surpassing 0.5 and RMSD below 3Å. The consistently high TMscores and low RMSD values also affirm the strong consistency between the generated backbones and their sketch inputs. This supports the confident classification of these generated backbones and their associated protein sketches as representing the same protein fold.

**Figure 3.**
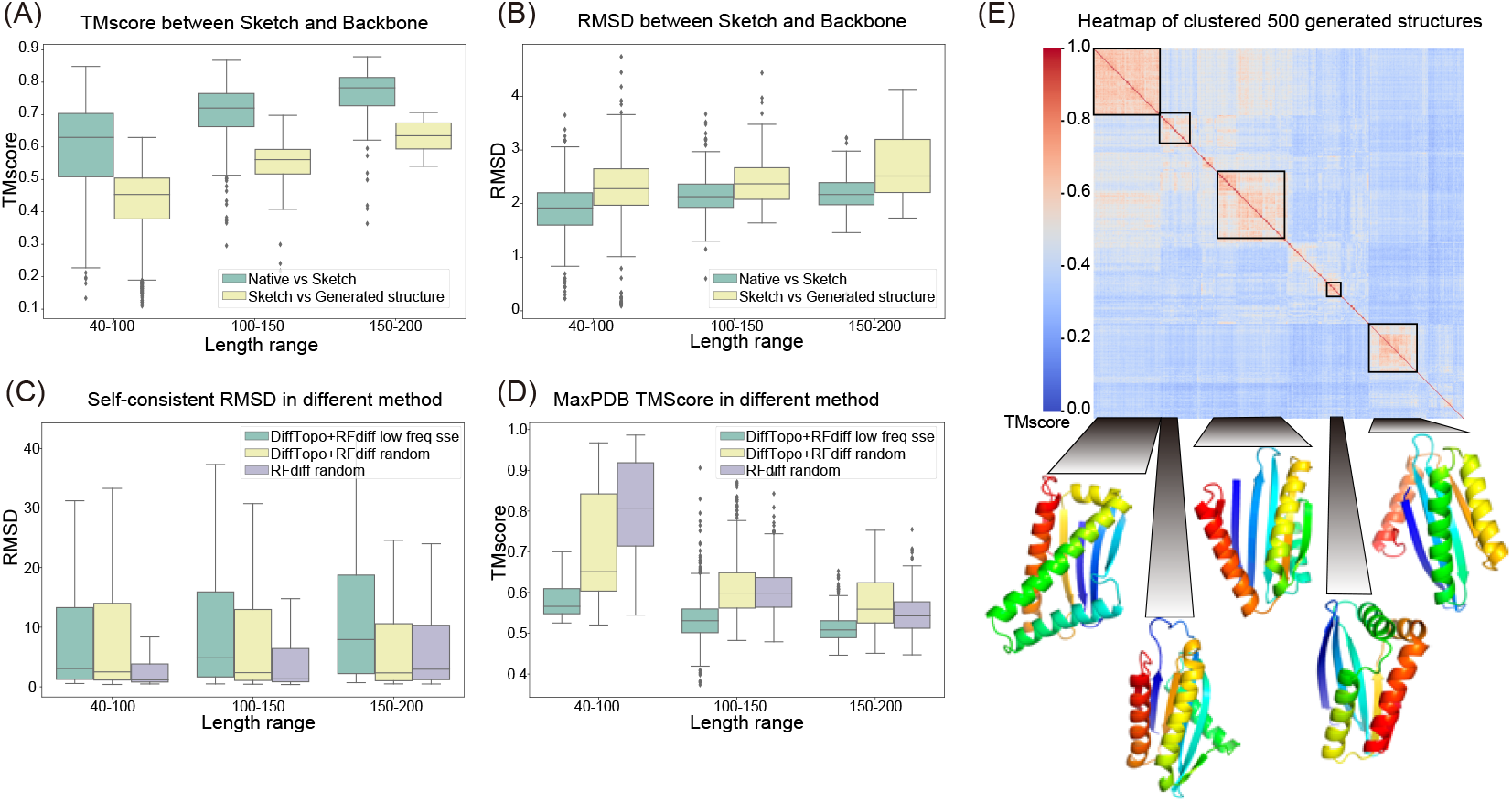
Characteristics of protein sketches and generated backbones: (A) RMSD between sketches and native structures (green) and generated backbones at different lengths (yellow). (B) TM-score between sketches and native structures (green) and generated backbones at various lengths (yellow). (C) Designability of generated backbones: scRMSD based on 1000 samples for each length range (40-100, 100-150, 150-200) in different methods. Purple: RFdiffusion random sample (baseline), Yellow: our pipeline’s random sampling of SSE strings, Green: our pipeline’s sampling of low-probability SSE strings in the database. (D) Novelty of generated backbones: color of box represents the same as in (C). (E) Diversity of generated backbones: heatmap of TM scores between each backbone after hierarchical clustering. Displaying structures from the top 5 largest clusters.

### 2.3 Designability, Novelty and Diversity

To test the designability of the backbone, it is necessary to design amino acid sequences for the backbone and predict whether these sequences can fold back into the corresponding topology. The designability of the backbone is evaluated through self-consistency assessment. Drawing inspiration from the research of FrameDiff (Yim et al., 2023b) and ProtDiff (Trippe et al., 2023), we quantify self-consistency using C*α* RMSD (scRMSD, lower values are better) and predicted local distance difference test (pLDDT, higher values are better). To assess the novelty of the backbone, we pick generated backbones with high designability (ScRMSD *<*3 Å and pLDDT *>*90) and search them in the entire CATH database and report their highest TMscore, referred to as Max PDB TMscore. To quantify diversity, we utilize the clustering function of FoldSeek (van Kempen et al., 2023) to cluster the sampled backbones with a 0.5 TMscore threshold. The TMscore between backbones and the number of clusters obtained are reported. We used RFdiffusion random sampling of backbones with lengths ranging from 40 to 200 as a comparative benchmark. Two types of random sampling were performed: random sampling of CG topology generated backbones using random lengths of SSE strings as input, and random sampling of CG topology using SSE strings with low occurrence frequency in the database. Figures 3C and 3D illustrate that, while there was a trade-off in designability compared to using RFdiffusion alone, numerous structures still maintained scRMSD values below 3Å. However, Figure 3D highlights that, when employing SSE strings with low frequency in the database for random sampling, our approach yielded significantly higher novelty in obtaining highly designable main chains compared to RFdiffusion. Figure 3E demonstrates the diversity in spatial arrangements of secondary structures sampled by our method. We sampled 500 highly designable backbones for the sequence ”EEEHHEHE” and calculated the TM score between each structure, resulting in a total of 109 clusters through clustering. Here, we present the representative structures of the 5 largest clusters.

## 3 Application for protein design

### 3.1 Novel fold exploration and Scaffold generation for functional motif

In order to showcase the model’s exploration capabilities across different folds, we selected three distinct secondary structure compositions: *α*, all-*β*, and mixed *α*-*β*, for protein fold exploration. Figure 4 illustrates the novel folds with high designability that we discovered in each of these secondary structure compositions. Our pipeline demonstrates remarkable versatility by uncovering novel folds that do not exist in various secondary structure combinations, including all-*α*, all-*β*, and mixed *α*-*β* folds. We also present the most similar structures to these folds that can be found with FoldSeek (van Kempen et al., 2023). Interestingly, these closest structures are not of the same fold as the ones we generated, demonstrating the novelty of our structures. Notably, even in the presence of 394 distinct 7-helix folds within the CATH database, our approach successfully identifies new and unique folds. By evaluating the sequence-to-structure quality of these main chains, we observe that the DiffTopo-RFdiffusion framework can sample novel folds and generate high-quality main chain structures.

**Figure 4.**
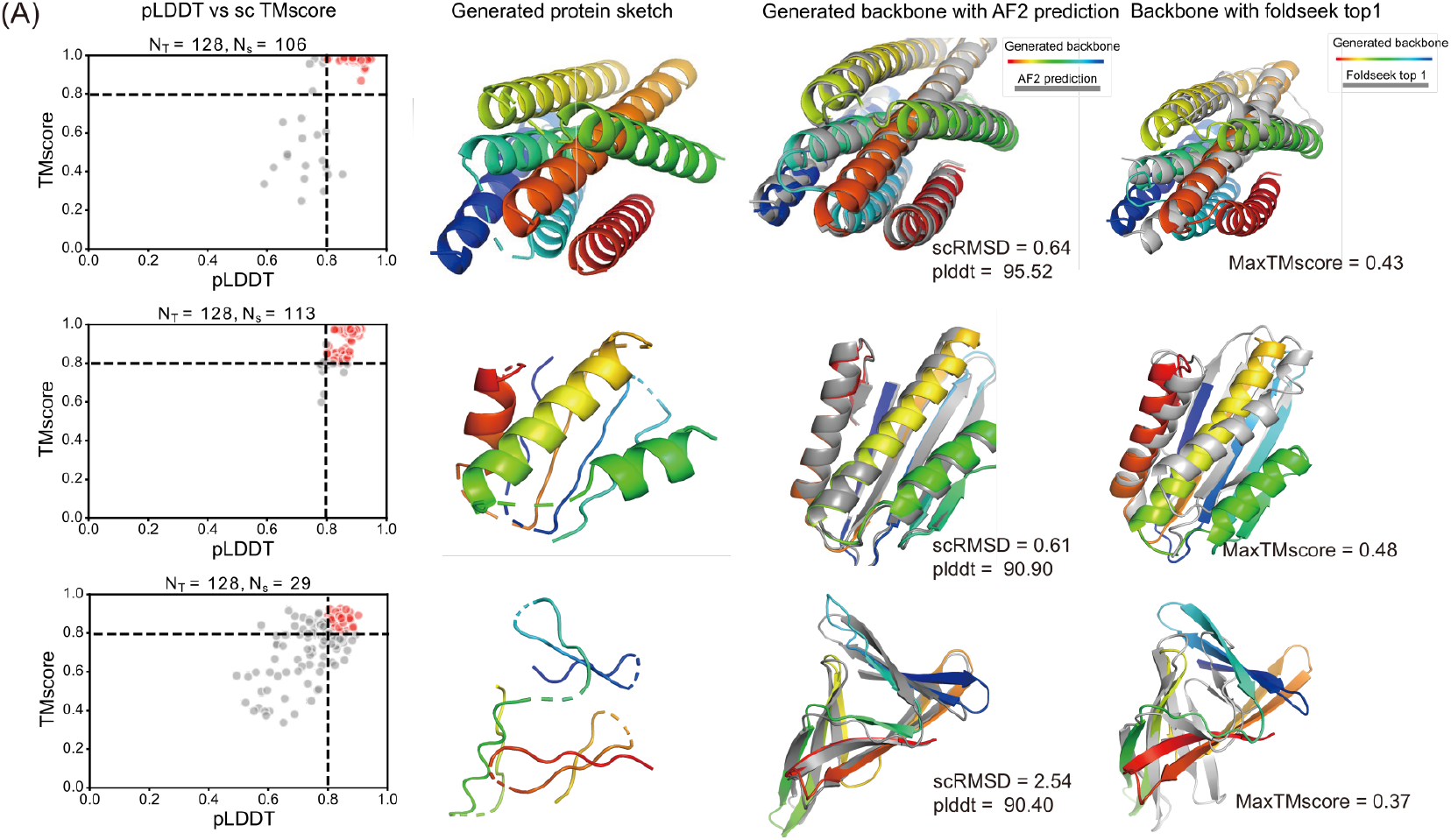
Computational design results for novel fold exploration in all-*α*(top), mixed *α*-*β*(middle) and all-*β* (bottom)folds. The plot shows TM-scores between generated backbones and Alphafold2 predictions against predicted plDDT. Red points indicate sequences with high designability (TM-score *>*0.8, plDDT *>*80). The second column displays DiffTopo-generated protein sketches. The third column presents backbone alignment with Alphafold2 predictions, and the last column compare the closest structure found by Foldseek.

In addition to random sampling of main chain structures, we explored another application of DiffTopo—generating scaffolds for existing secondary structure motifs. Here, we attempted to design a new scaffold for the DBL1 03 protein binder that binds to PD-L1 (complex PDB id: 7XYQ) (Gainza et al., 2023). The A1-A20 helical portion of DBL1 03 forms the interacting interface with PD-L1. We developed a motif scaffolding pipeline (in Appendix A.1.4). The best-designed structure, predicted using Alphafold multimer, is shown in the right panel of Figure B.1 in the Appendix. It is evident that the predicted binding positions by Alphafold align well with the ground truth. This demonstrates the potential of our method in constructing scaffolds for tasks involving the creation of motifs with existing secondary structures.

## 4 Discussion

We show that a new representation termed CG topology is able to represent protein folds correctly using very few data points, reducing the sampling space for protein design. With our generative diffusion model, DiffTopo, we not only learn the distribution of relative positions of secondary structures but also sample diverse topologies. By passing protein sketches to RFdiffusion, we are able to generate designable protein backbones in 3D space and explore the protein fold space with fewer degrees of freedom. Comparing with RFdiffusion only as benchmark, and using ProteinMPNN and AlphaFold2 as an orthogonal test, our framework improves the diversity of protein space exploration while still maintaining good designability. Our model also shows the capability in motif scaffolding problem when motifs with secondary structures are available.

Our results demonstrate that the DiffTopo+RFdiffusion framework is capable of designing new proteins and explore novel folds non-existent in nature. Compared to all other backbone atom level models like RFdiffusion and Framediff, our framework can massively sample topological space with different secondary structure arrangements in a short time. On the other hand, compared to Form and GENESIS(Harteveld et al., 2022), we can make protein topology sampling automatically instead of manually construct it. Our work provides a new perspective on protein structure representation and we believe there is much to explore in the space of this representation to improve performance in protein universe searching and protein design. For example, our method can be leveraged to generate customized protein backbones exhibiting highly regular patterns, similar to nanomaterials. Moreover, our approach holds the potential to create novel scaffolds for existing functional motifs, offering applications in de novo enzyme and antibody design.

## A Appendix

### A.1 Method

#### A.1.1 Coarse-grained topology representation

Inspired by the Form representation (Taylor et al., 2008) and protein sketches (Harteveld et al., 2022), where protein structures are built by the assembly of their secondary structures, we adopt a simplified representation referred to as a ”Coarse-grained topology (CG topology)”. The topology reduces the intricate protein structure into a stacked arrangement of Secondary Structure Elements (SSEs), achieved by representing secondary structures with three carbon alpha centroids, as illustrated in Figure 1A. CG topology involves capturing the geometric position of helices and strands through centroids. For helices, centroids include the first and last four C*α* atoms, as well as the total C*α* atom centroid. Strands are represented by centroids of the first and last two C*α* atoms, along with the overall C*α* atom centroid within the strand. CG topology can also be easily represented as a protein sketch, which is a rough 3D approximation of a native protein structure with standard SSEs, lacking loops, and AA side chains. Compared with Form and protein sketch, this representation method of CGtopo gets rid of the concept of layers and has a higher freedom in structural representation to represent structures like beta barrels, and it is still simple with a small number of degrees of freedom.

#### A.1.2 Dataset

Using the CG topology approach we can easily convert a protein structure database, such as PDB, from standard protein models to CG topology structures. In this paper, all PDB structural data are sourced from CATH, a database organized as a classification of protein structures (Knudsen & Wiuf, 2010). To mitigate the influence of structural redundancy on the data distribution, we employ the CATH-dataset-nonredundant S40 for training data, yielding 31,886 non-redundant protein structures. 90% of structures are used for training and 10% rest are used for independent validation.

#### A.1.3 DiffTopo+RF diffusion F ramework

First, we introduce DiffTopo, a Equivariant diffusion model for generating CG topology conditioned on SSEs strings. Based on previous works on denoising diffusion (Song et al., 2021; Ho et al., 2020; Austin et al., 2023), given a data point sampled from a real data distribution *z*_0_*∼ q*(**z**), the forward diffusion process is defined by adding Gaussian noise gradually to the data point *z*_0_ form corrupted samples *z*_*t*_. For one CG topology data *z*_0_, there are three points *x* and corresponding secondary structure type *u*. At time step *t* = 0, …, T, the conditional distribution of the intermediate data state *z*_*t*_ given the previous state is defined by the multivariate normal distribution,

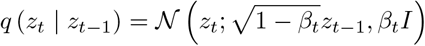

The step sizes are controlled by a variance schedule 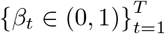. The process is constructed to be Markovian and if we let *α*_*t*_ = 1 − *β*_*t*_ and 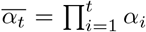 we could obtain the distribution of *z*_*t*_ given *x*

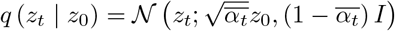

For the backward process, we need to learn a model *p* to approximate these conditional probabilities.

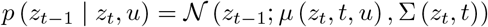

*µ*_*θ*_ (*x*_*t*_, *t, u*) is predicted by the neural network. In this paper function *µ* that predicts noise *ϵ*_*θ*_ is implemented as geometric vector perceptrons (Jing et al., 2021). The input to GVPs is the noised version of the point coordinates *x*_*t*_ and point feature *h*_*t*_ at time *t* and context *u*. Note that the predicted noise *ϵ* includes coordinate and feature components, *ϵ* = [*ϵ*_**x**_, *ϵ*_**h**_]. The predicted noise could be calculated by

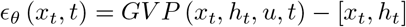

Following the generation of the CG topology via the diffusion process, utilizing CG topology allows us to straightforwardly obtain the positions and lengths of standard Secondary Structure Elements (SSEs). Since CG topology retains a certain level of fuzziness, we need a backbone level model to systematically search for the designable backbone corresponding to CG topology. Here, we leverage RFdiffusion, a highly effective backbone generation model to generate designable backbones from CG topology. Initially, we construct a protein sketch from standard SSEs based on CG topology and utilize the motif modeling function inherent in RFdiffusion to directly connect the SSEs. Subsequently, we employ the partial diffusion approach to identify a reasonable backbone structure within this interconnected structure. Then we employ the Protein Message Passing Neural Network (ProteinMPNN) (Dauparas et al., 2022) for fixed backbone sequence design algorithm to obtain amino acid sequences. These sequences are then input into the structure prediction algorithm Alphafold2 (Jumper et al., 2021) to predict the structure, which is use to validate the designability of generated backbones.

#### A.1.4 motif sccafold modeling

First, we extract the initial helix combined with PD-L1 in DBL 03 and convert it to CGtopo. Subsequently, we employ Difftopo’s conditional generation protocol to sample the entire scaffold’s CG- topo, using the condition ’HEHEHE’ while keeping the first helix unchanged. The sampled CGtopo is then constructed into a protein sketch, with the first helix replaced by a functional motif.

Next, RFdiffusion’s fixcontig protocol is utilized to connect the loops, followed by the partial diffusion protocol to optimize the entire backbone while preserving the first 20 amino acids. Lastly, ProteinMPNN is employed to design sequences for positions other than the first 20 amino acids. Alphafold is then used to predict the monomer structure, and Alphafold multimer is applied to predict the complex structure.

## B Supplementary figures

**Figure B.1.**
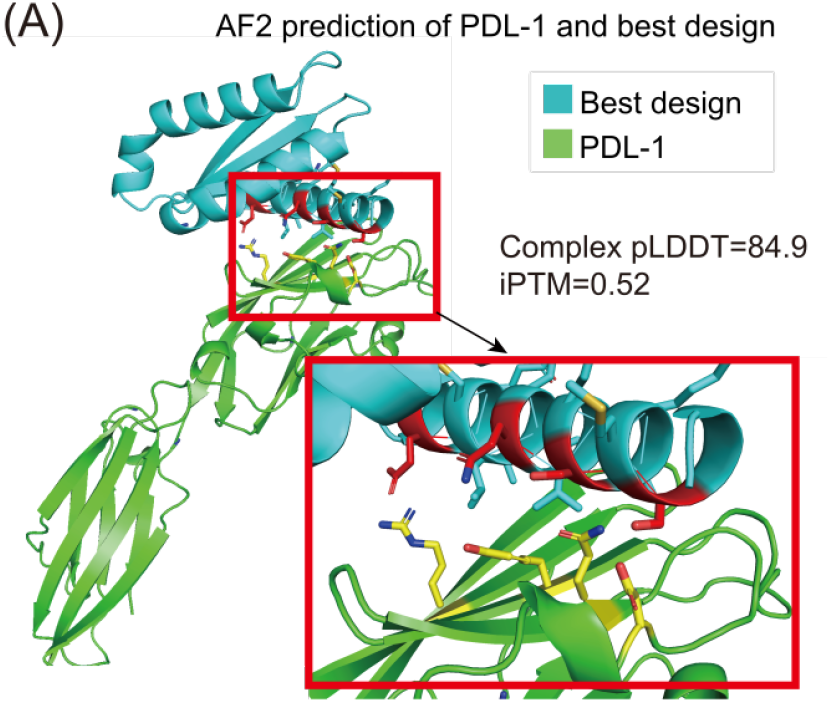
Motif scaffolding application. (A) Prediction of multimer structure of best design and PD-L1. Zoomed-in part shows the AlphaFold2-predicted interface, with red residues representing the functional motif binding to PD-L1.

